# Radiomics Analysis Using Stability Selection Supervised Principal Component Analysis for Right-censored Survival Data

**DOI:** 10.1101/408831

**Authors:** Kang K. Yan, Xiaofei Wang, Wendy Lam, Varut Vardhanabhuti, Anne W.M. Lee, Herbert Pang

## Abstract

Radiomics is a newly emerging field that involves the extraction of a large number of quantitative features from biomedical images through the use of data-characterization algorithms. Radiomics provides a noninvasive approach for personalized therapy decision by identifying distinctive imaging features for predicting prognosis and therapeutic response. So far, many of the published radiomics studies utilize existing out of the box algorithms to identify the prognostic markers from biomedical images that are not specific to radiomics data. T o better utilize biomedical image, we propose a novel machine learning approach, stability selection supervised principal component analysis (SSSuperPCA) that identify a set of stable features from radiomics big data coupled with dimension reduction for right censored survival outcomes. In this paper, we describe stability selection supervised principal component analysis for radiomics data with right-censored survival outcomes. The proposed approach allows us to identify a set of stable features that are highly associated with the survival outcomes, control the per-family error rate, and predict the survival in a simple yet meaningful manner. We evaluate the performance of SSSuperPCA using simulations and real data sets for non-small cell lung cancer and head and neck cancer, and compare it with other machine learning algorithms. The results demonstrate that our method has a competitive edge over other existing methods in identifying the prognostic markers from biomedical big imaging data for the prediction of right-censored survival outcomes. An R package SSSuperPCA is available at the website: http://web.hku.hk/∼herbpang/SSSuperPCA.html

## I. INTRODUCTION

Bioimaging has emerged as an important diagnostic and prognostic tool in cancer. Imaging, as a non-invasive technique, has an advantage over other invasive clinical options and can generate a large amount of radiomics texture features that might be useful for predicting prognosis and therapeutic response for various conditions. In addition, it can provide valuable information for individualized treatments. Previous studies for analyzing bioimaging data have mostly focused on categorical outcomes. Success stories include the diagnosis of brain tumors through sparse representation-based quantitative method on magnetic resonance imaging (MRI) [1], the forecast of non-small cell lung cancer nodule recurrence by leveraging the texture information contained in radiology imaging such CT [2], and the prediction of the progression of adolescent idiopathic scoliosis with 3-D spine biplanar X-ray images [3].

Radiomics texture features are of high dimension in nature, therefore traditional statistical and computational models are not suitable. Recently Cheng et al. introduced machine learning classification methods for radiomics biomarkers to predict survival by dichotomizing the censored continuous survival data at a specific time cutoff [4]. The objective is to stratify patients into two survival classes by relevant survival time. However, due to the right censored nature of the data, dichotomizing the survival time into binary outcome will lead to loss of information. A recent effort has been conducted to evaluate existing prognostic modeling methods for radiomics data with 1,610 features [5]. The researchers found that the combinations of machine learning algorithms and feature selection methods would have an impact on the prediction of overall survival. In their paper, the highest median performance of predicting overall survival was shown by the combination of either boosted gradient Cox model or survival regression with four feature selection methods that including mutual information maximization, minimum redundancy maximum relevance, univariate Cox regression or no feature selection. However, none of these combinations stands out in their study as their performances were quite similar when applied to one real data set.

For radiomics study, it is important to assess the clinical relevance of radiomics features and examine its performance and stability for predicting prognosis. Biomarker predictors obtained from one-time experiment may not be easily generalizable. Also, texture features with higher stability tend to be more informative and have higher prognostic performance as well as reproducibility. This is demonstrated by Aerts et al. [6]. To the best of our knowledge, many of the radiomics studies were conducted by using some existing out of the box algorithms that are not specific to radiomics data. Unlike genomics [7], currently there is a lack of prognostic algorithms that are specifically designed for high-dimensional radiomics data. In this study, we focus on modeling texture features for cancer prognosis prediction with over 10,000 features extracted from biomedical images.

Here we present a novel algorithm SSSuperPCA, a stability selection supervised principal component analysis tool for radiomics data, and apply it to two real radiomics data sets. Our objective is to address the issue of the lack of novel prognostic tools that directly predict right censored data for radiomics texture big data. We also benchmark our proposed SSSuperPCA against other regression and machine learning methods.

## II. MATERIALS AND METHODS

There are two major components involved in the proposed algorithm stability selection supervised principal component analysis (SSSuperPCA). Our proposed algorithm - stability selection supervised principal component analysis - integrates stability selection with supervised principal component analysis for right-censored survival data.

Stability selection was first introduced by Meinshausen and Buhlmann [8]. It is a generic approach that can be applied to a wide range of statistical techniques for feature selection in high-dimensional data. In contrast to other feature selection algorithms that aim to find the best predictors, stability selection identifies a set of stable features that are chosen with high probability with the rationale that they can provide consistent predictions. The idea of stability selection is to perform a feature selection algorithm on many subsets with observation and features that are randomly resampling from the original data. Selected features will be aggregated after repeating this procedure for a number of times. Average selection probabilities will be computed for each feature, and we can expect strong relevant features will have higher selection probabilities close to 1 while irrelevant features will have probabilities close to 0 [8]. Shah and Samworth [9] proposed a refined version of stability selection called complementary pairs stability selection that uses the subsample as well as its complementary pairs. A narrower error bound could be derived with the modified Markov’s inequality that assumes unimodality or r-concavity for the distribution of the selection frequencies. In order to tackle high-dimensional feature selection for-omics data with right-censored survival outcomes, Mayr et al. (2016) proposed a boosted C-index stability selection that combined complementary pairs stability selection and C-index boosting to filter out informative features and to acquire the risk prediction that maximizes its discriminatory power [10]. Bair et al. first proposed the supervised principal components analysis that integrates principal component analysis (PCA), an orthogonal linear transformation technique for dimensionality reduction, with generalized regression to address the issue of high-dimensional data [11]. When dealing with number of features far exceeding the number of observations, the conventional method may yield unsatisfactory results because of sparsity. Supervised PCA can generate favorably results through the use of the subsets of the selected features that account for their correlation with the outcome. This approach can reduce the potential problems caused by the noisy features on the prediction model and keep the model’s simplicity at the same time. Supervised PCA has been applied to the area of bioinformatics studies for most scenarios, such as genome-wide association analysis [12] and microarray gene expression analysis [13]. Radiomics, the recently emerging research field which extracts high dimensional quantitative features from medical images by using advanced data-characterization extraction algorithms, could also benefit from supervised PCA. Kickingereder et al. first incorporated supervised PCA with radiomics to provide a noninvasive approach as a novel decision tool for improving decision-support in the treatment of glioblastoma patients [14].

The analysis was performed using our R package, SSSuperPCA that was built on mboost, stabs and superpc under R version 3.4.3.

### A. Stability Selection Supervised Principal Component Analysis (SSSuperPCA)

SSSuperPCA identifies the informative features from a large set of quantitative features extracted from biomedical images and predicts right-censored survival outcomes with supervised PCA to obtain the prediction model. Here, the continuous additive predictor *η* derived from the prediction model is defined as follows:

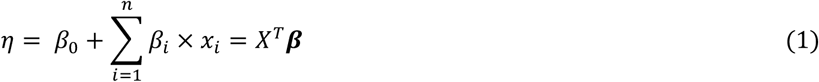

where ***X = (x*_*1*_, *x*_*2*_,** …, ***x*_*n*_)^*T*^** is the feature vector and **β**=(β_1_,β_2_…β_n_**)^*T*^** is the corresponding generalized linear regression coefficients vector.

This algorithm has seven basic steps:

1. **Categorize** the features into *M* subgroups based on the decreasing order of the average Uno’s truncated C-index that was derived from the fc-fold cross-validation of a univariate Cox proportional hazards regression model;
2. **Apply** the complementary pairs stability selection with boosted C-index to the 2 X B subsamples for each subgroup.
  a. **Initialize** the estimation of the continuous additive predictor 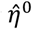 for each subsample and the corresponding features in each subgroup. For example, set 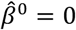 which will make 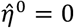. Set ***m = 1*** and a large maximum number of iterations ***m***_stop_.
  b. **Compute** the negative gradient vector of the loss function and figure out its value at 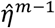 that was derived from the previous iteration.
  c. **Fit** the negative gradient vector of the loss function to ***x*_*t*_,** the features in the subgroup, through the base learners Cox proportional hazard regression model *b*_*1*_(·).
  d. **Select** the feature that minimizes the value of the loss function under the least squares criterion.
  e. **Update** the continuous additive predictor with the feature selected in d).
  f. **Stop** if ***m*** meets the stop value ***m*_*stop*_.** Otherwise update ***m = m + 1*** and return to step a).
  g. **Average** the selection probabilities that are derived from the 2 × B subsamples and return features that exceed the preset threshold
3. **Aggregate** the selected features obtained from each subgroup.
4. **Estimate** the standardized correlation coefficients, like proportional hazards for survival data, for each feature selected form stability selection through the generalized univariate linear regression.
5. **Construct** the reduced features matrix by using features whose absolute correlation coefficients exceeds the threshold that estimated via cross-validation.
6. **Compute** the principal components of the reduced features matrix.
7. **Predict** the right censored survival outcome in the Cox proportional hazards regression model by using the first few principal components and return the stable features selected in step 3).

Unlike the conventional principal component analysis, SSSuperPCA performs singular value decomposition on the reduced data matrix to incorporate features that are highly correlated with the outcome. The input feature matrix for SSSuperPCA in step 4, whose components are features resulted from the stability selection with boosted C-index, should be standardized before performing prediction.

#### 1) Stability Selection with Boosted C-Index

The concordance index (C-index) is a routine criterion in biomedical studies that measures the rank-based agreement probability between the continuous additive predictor and right censored survival outcome. C-index may take values from 0.5(random predictor) to 1 (perfect prediction accuracy).

In general, C-index provides a global assessment of the discriminative ability of the fitted survival models. However, C-index may result in biased estimation because of the censoring pattern due to the fact that the events are not observable for all patients in practice. Observation pairs that cannot be ordered due to censoring will be omitted in the evaluation. T o overcome this shortcoming, Uno proposed the truncated C-index that is independent with the censoring distribution and could consistently produce an asymptotically unbiased estimation of the conventional C-index [15]. The truncated C-index estimator is formulated as below:

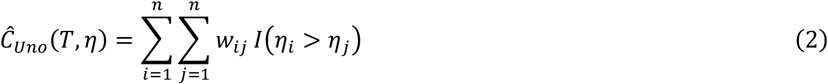

Where 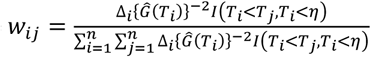, with ***T*** denotes the observed survival time subject to censoring, G(.) is the Kaplan-Meier estimator of the survival function that accounts for the censoring time, Δ is the censoring indicator and ***I*** (.) is the indicator function.

To obtain the optimal generalized linear regression coefficient *β* that maximize the truncated C-index, the component-wise gradient boosting algorithm [16] which is computationally efficient for high-dimensional data has been adopted. The idea is to update the base-learner in each boosting iteration through adding features that best fit the least square criterion of the negative gradient vector that derived from the loss function. However, Uno’s truncated C-index estimator it is non-differentiable with respect to ***r],*** directly using it as the loss function for gradient boosting is out of the question. Therefore, an approximate approach that uses the sigmoid function has been applied to smooth the Uno’s truncated C-index [17].

The smoothed estimator is given as below.

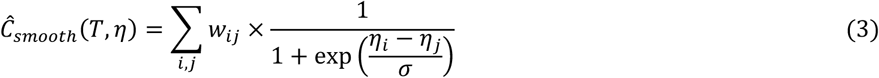

σ is a positive parameter which controls the accuracy of the sigmoid approximation. Previous studies demonstrated that the sigmoid approximation and its corresponding outcomes were not sensitive to σ when σ is small enough [17].

To avoid the problems of overfitting while ensuring the selection of informative features, boosted C-index was incorporated with complementary pairs stability selection proposed by Shah and Samworth. The general concept is to use 2 × B complementary pairs of size n/2 (the subsamples and its complement as well) and fit the boosted C-index model to select a pre-specified number of features on each of the subsamples. Average selection probabilities will be calculated for each feature after aggregating the selection results of all the **2 × B** subsamples and only features that beyond a threshold will remain in the final selection list. By cooperating with stability selection, we can easily control the upper bound of per-family error rate (PFER) with the inequality showed below.

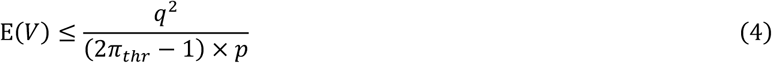

where E(*V*) is the expected number oi false positive selected features, *π*_***thr***_ is the probability threshold for stability selection, ***q*** is the number of features that need to be selected on each subsample, and *p* is the number of total features.

PFER metric, as the main characteristic of stability selection, can strictly control the noisy features that are falsely included. Meanwhile, the features matrix extracted from the biomedical images could be sparse (the number of features is much greater than the number of observations) or the features are not highly correlated with the outcomes. Thus, less features would be selected when applying the stability selection with boosted C-index algorithm to radiomics data. To tackle this problem and to include more informative features as the input for Supervised PCA. We first grade all the features based on the Uno’s truncated C-index that derived from a univariate generalized linear regression model and categorize the features into subgroups based on the ranking, then we perform stability selection with boosted C-index on the subgroups and aggregate the features selected from each subgroup as the final conclusion. In our experiments, the number of complementary pairs (B) was set to 50 which led to 100 subsamples, and the number of uniquely selected variables ***(q)*** in each subsample was set to an upper bound as a function of B and the number of total features (*p*). The probability threshold for stability selection ***n*_*thr*_** was chosen to be slightly higher than 0.5 to include more features selected from biomedical image and to reduce possible information loss.

### 2) Supervised PCA

The features that are selected through the stability selection with boosted C-index will serve as the entry matrix for Supervised PCA. Denote *X*_*n × p*_ as the standardized feature matrix, where ***n*** is the number of observations and p is the number of features.

First, the standardized regression coefficients *C = (β*_*1*_, *β*_*2*_ …, *β*_*p*_) that measure the effect on the survival outcome will be estimated through the univariate Cox proportional hazards regression model. Let 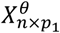 be the new feature matrix that consists the columns of *X*_*n × p*_ whose absolute value of the standardized regression coefficients exceed the threshold *θ*. Here the optimal value of *θ* is determined by cross-validation of the log partial-likelihood ratio statistics. The singular value decomposition of 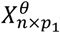 is 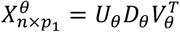, where the dimension of *U*_θ_, *D*_θ_, and *V*_θ_ are *n × m,m × m,p*_*1*_*1 × m*, and *m* ***=*** min(*n,p*_*1*_). 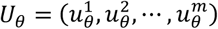 is the supervised principal component of 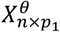. The last step is to use the first few supervised principal components to fit the Cox proportional hazards regression model and the number of supervised principal components used in the prediction is less than three in general. The threshold of regression coefficient *θ* and the number of supervised principal components used in our experiments were determined through 5-fold cross-validation.

#### 3) Comparison with Other Machine Learning Algorithms

We compared the stability selection supervised principal component analysis and different machine learning algorithms for radiomics survival data. The algorithms compared including supervised principal component analysis without stability selection, Cox proportional hazards regression model [18], lasso and elastic-net based Cox proportional hazards regression model [19], and survival random forests [20]. The basic cox proportional hazards regression is implemented with R package survival, and the univariate Cox regression was employed to filter out the top ten statistically significant features, then the forward and backward stepwise Cox regression were applied to build the prediction model. The lasso and elastic-net based Cox regression approaches are available as Coxnet in R. 5-fold cross-validation was used to determine the optimal shrinkage parameter X for lasso and elastic-net based Cox regression. The weight for L1 penalty and L2 penalty terms in elastic-net Cox regression is fixed to 0.5. The survival random forests is carried through R package randomForestSRC, with the number of trees equals to 1000 and the split rule is set to ‘log-rank’.

### B. Data Sets and Generation of Texture Features

We considered the two data sets, head and neck cancer [21] and non-small cell lung cancer [22], which are available on The Cancer Imaging Archive (TCIA) [23]. Only Computed Tomography (CT) and its corresponding RTSTRUCT biomedical image files in both data sets are used for data extraction procedure. The feature extraction part was conducted with Matlab by using the algorithm proposed by Vallieres et al. [24]. There are total 4 non-texture features and 43 texture features extracted from the tumor regions. The non-texture features are volume, size, solidity, and eccentricity, and the texture features include gray-level co-occurrence matrix (GLCM), gray-level run-length matrix (GLRLM), gray-level size zone matrix (GLSZM) and neighbourhood gray-tone difference matrix (NGTDM). Details of these features could be found in the original paper. Different extraction parameters setting were evaluated to investigate the influence of the extraction parameters on the predictive texture features. The values for wavelet band-pass filtering, isotropic voxel size, and quantization of gray level have followed the suggestion of original paper. General information regarding sample size and the number of features of the two data sets are listed in Table 1.

**Table 1.**
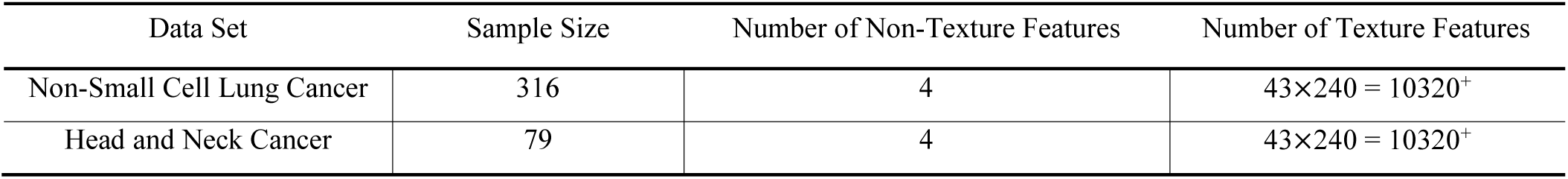
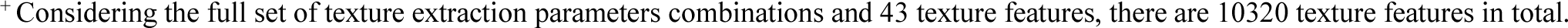
Radiomics data sets used.

| Data Set | Sample Size | Number of Non-Texture Features | Number of Texture Features |
| --- | --- | --- | --- |
| Non-Small Cell Lung Cancer | 316 | 4 | $43 \times 240 = 10320^+$ |
| Head and Neck Cancer | 79 | 4 | $43 \times 240 = 10320^+$ |

<sup>+</sup> Considering the full set of texture extraction parameters combinations and 43 texture features, there are 10320 texture features in total.

The 10,324 features extracted from the CT/RTSTRUCT image files are used to predict the overall survival of the patients, and a risk score will be applied to categorize the patients into high-risk and low-risk groups. The measurement matrices used in our experiment are Uno’s C-index that assesses the discriminative ability of the model, Brier score that indicates the overall model performance and the log-rank statistics that evaluate whether the survival time of the two groups is statistically significantly different or not.

In order to understand how well our approach performs under different situations, we also performed simulations with the non-small cell lung cancer data. With the stable features set obtained from our 100 times experiments, we randomly drew 3,000 features from the original data. Some of the 3,000 features were selected from a stable feature set while others were chosen from less informative features with stability selection probability equal to 0. Under the alternative, we simulated nine situations under different sample sizes with different number of stable features. The sample size is set to 150, 200 and 250 while the number of stable features is assigned to 5, 15 and 25 respectively. In addition, we simulated the null cases with 3000 features by capturing the mean and covariance matrix from features with C-index around 0.50 (plus or minus 0.02), and the sample sizes were generated as above. In total, we simulated twelve different scenarios.

## III. RESULTS

In this section, we applied our algorithm to assess its abilities in predicting the survival from biomedical images of the two cancer data and compared its performance with the five machine learning algorithms. For the two data sets, we randomly selected 65 percent of the observations as a training set to build the model, and the remaining 35 percent was used as a testing set to evaluate the performance of the models. This procedure was repeated for 100 times to calculate the average values for the three measurements mentioned above.

### A. Simulation Studies

We simulated 100 data sets for each of the situations with different sample size and the number of stable features as described in section 2.4. For each data set, we split it into training and testing set and estimated the corresponding measures of Uno’s C-index, Brier score, and log-rank statistics.

The detailed results of Uno’s C-index for alternative scenarios are shown in Table 2, and the remaining two measures of alternative cases, as well as the results for null scenarios are provided in the supplementary Tables 1-3. For the results of null scenarios, the Uno’s C-index for different sample size were 0.492, 0.491 and 0.511, respectively. Considering the results of the alternative scenarios, as the number of stable features increased from 5 to 25 along with the number of sample size increased from 150 to 250, the Uno’s C-index increased from 0.550 to 0.624, the Brier score dropped from 0.152 to 0.148 and the log-rank statistics increased from 2.914 to 6.104. We clearly observed that as the number of top stable features increases, the performance of SSSuperPCA improves. The same pattern could also be identified for the sample size, while the impact was not as large as the number of top stable features. However, as the sample size and the number of stable features increased, the improvement of its performance became less impressive.

**Table 2.** Average Uno’s C-Index of the simulated data sets under different scenarios

| Number of informative features over 3000 features | Size 150 | Size 200 | Size 250 |
| --- | --- | --- | --- |
| 5 | 0.550 | 0.552 | 0.561 |
| 15 | 0.586 | 0.586 | 0.589 |
| 25 | 0.613 | 0.621 | 0.624 |

### B. Application to non-small cell lung cancer data set

The non-small cell lung cancer (NSCLC) data contained 208 events out of 316 observations and the median survival time was 543 days. The results of non-small cell lung cancer data showed that our proposed algorithms SSSuperPCA, which integrate stability selection with boosted C-index and supervised PCA, performed better than supervised PCA without stability selection. SSSuperPCA increased the Uno’s C-index from 0.588 to 0.628, and the log-rank statistics from 5.654 to 7.794 when comparedwith supervised PCA without stability selection. The Brier score of SSSuperPCA was lower than that of supervised PCA, which suggested that prediction model of SSSuperPCA was also better. We also aggregated the stable features that selected through our algorithm SSSuperPCA in the 100 times’ running to further understanding the stable features that contribute most to predict the survival of non-small cell lung cancer patients. In summary, the top three features were selected as the stable features for more than 90 times. The highly selected features including two non-texture features, volume, size and one texture features of small zone emphasis. Volume and size are two features that describe the size and shape of the tumor region and small zone emphasis is to assess homogeneous of the texture. It is acceptable that the tumor region characteristic would have higher prognostic power for survival prediction of non-small cell lung cancer patients. Detailed definition of these features could be found in Vallieres’s paper.

Table 3 presents the average measurements for the six algorithms with non-small cell lung cancer data. In general, the performance of SSSuperPCA was higher than other algorithms compared using this data set based on the Uno’s C-index, Brier score, and log-rank statistics. For this data set, we observed that the survival random forests generated the worst prediction for survival, only slighter better than a random guess. For Cox proportional hazards regression model, lasso and elastic-net based Cox proportional hazards regression model, these algorithms produced quite similar results with Uno’s C-index around 0.590, Brier Score equal to 0.152, and log-rank statistics close to 4.50. Figure 1 shows the scatter plots of the SSSuperPCA against the other four algorithms when considering Uno’s C-Index (scatter plots of the SSSuperPCA against Cox model are provided in Supplementary Figure 1). Points above/on the reference line means that Uno’s C-index of SSSuperPCA is no less than the value of algorithms compared. The scatter plots for Brier score and log-rank statistics are provided in Supplementary Figure 2.

**Table 3.**
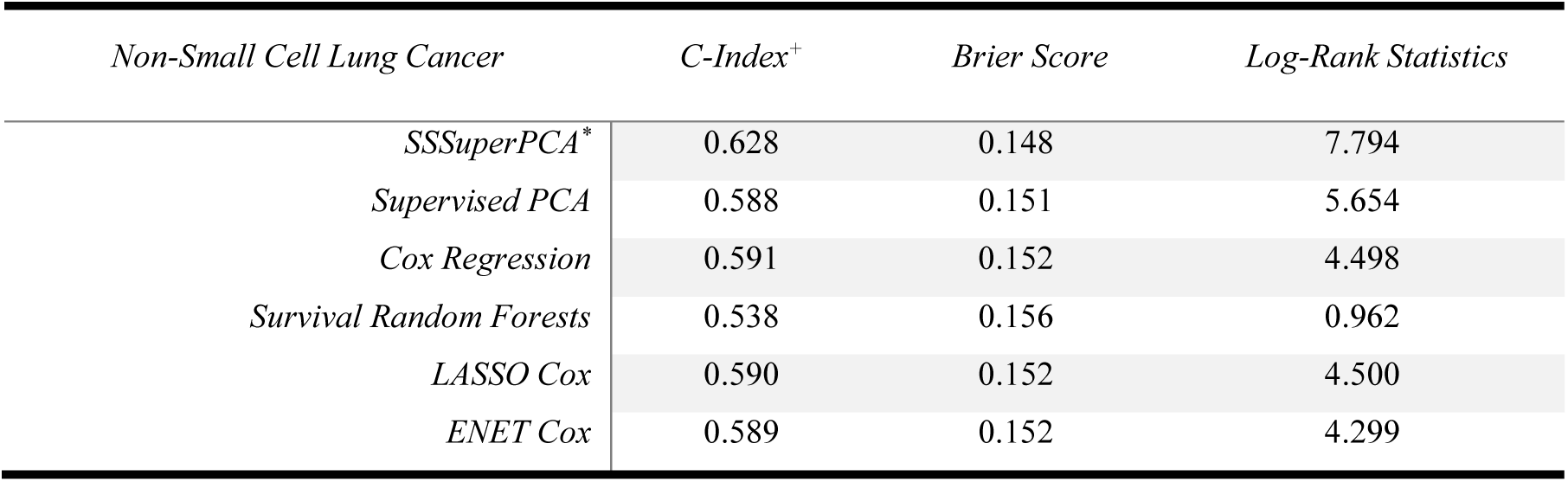
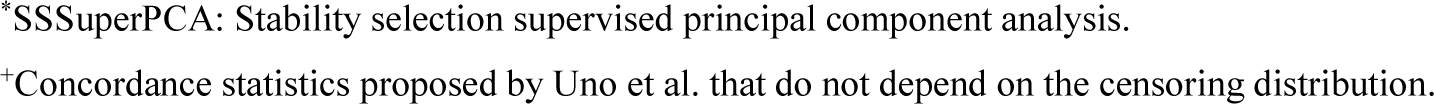
The comparison of stability selection supervised principal component analysis with five machine learning algorithms with data nonsmall cell lung cancer. The results are the average values that based on 100 times random training/testing splits.

| *Non-Small Cell Lung Cancer* |  | *C-Index<sup>+</sup>* | *Brier Score* | *Log-Rank Statistics* |
| --- | --- | --- | --- | --- |
|  | *SSSuperPCA\** | 0.628 | 0.148 | 7.794 |
|  | *Supervised PCA* | 0.588 | 0.151 | 5.654 |
|  | *Cox Regression* | 0.591 | 0.152 | 4.498 |
|  | *Survival Random Forests* | 0.538 | 0.156 | 0.962 |
|  | *LASSO Cox* | 0.590 | 0.152 | 4.500 |
|  | *ENET Cox* | 0.589 | 0.152 | 4.299 |

\*SSSuperPCA: Stability selection supervised principal component analysis.
<sup>+</sup>Concordance statistics proposed by Uno et al. that do not depend on the censoring distribution.

**Figure 1.**
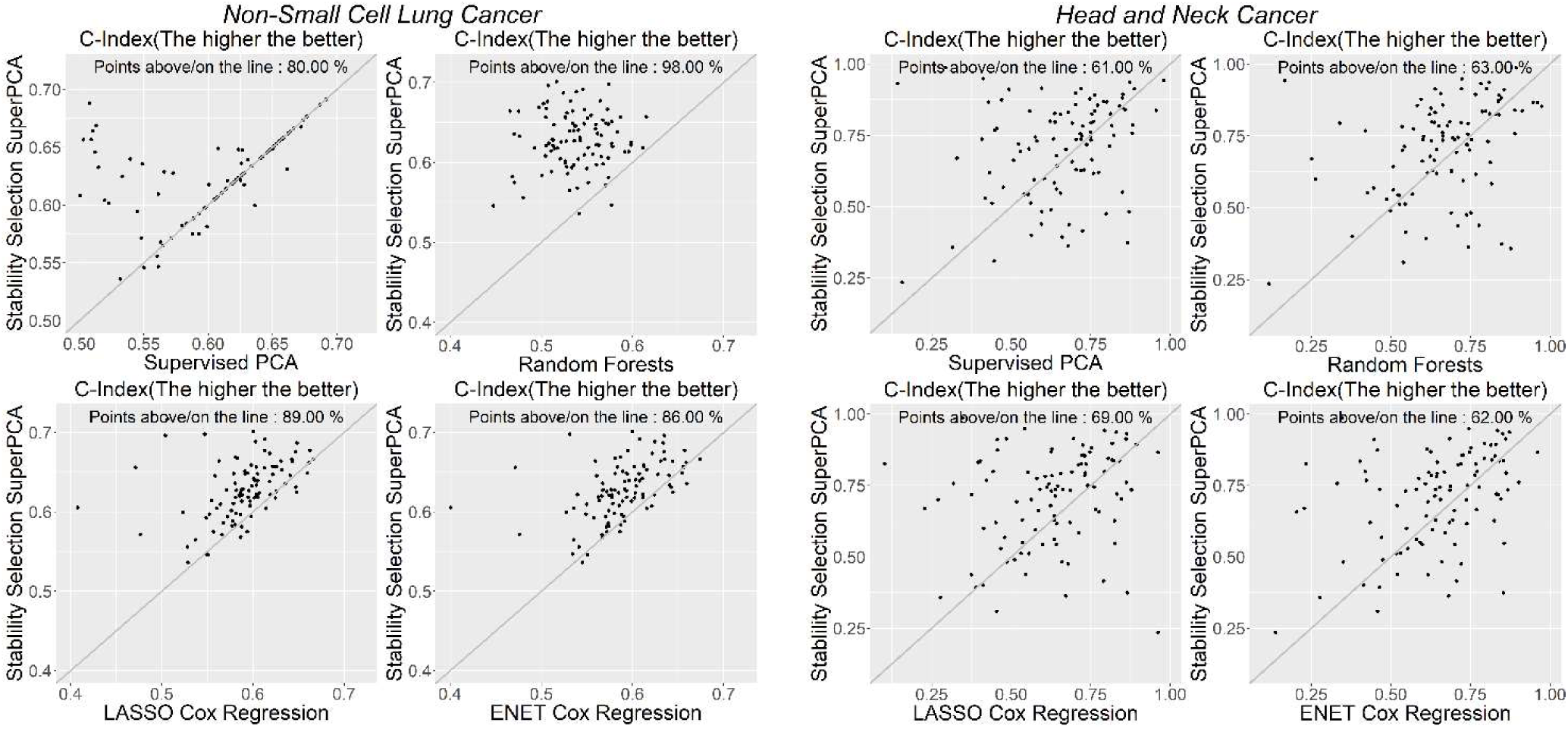
Scatterplots of Uno’s C-index for non-small cell lung cancer data.

In additional, we tested the top machine learning algorithms combined with feature selection methods that were mentioned in Leger’s study [5]. For non-small cell lung cancer data, the boosted gradient linear model combined with univariate selection could achieve the best performance, with Uno’s C-index, Brier score, and log-rank statistics equal to 0.589, 0.153, and 4.074, respectively.

### C. Application to head and neck cancer data set

The patients of head and neck cancer data set we used were collected from the two cohorts, 30 observations from Centre hospitalier universitaire de Sherbrooke and 49 observations from Hopital general juif de Montreal, QC, Canada. Before performing survival analysis, a log-rank test was used to evaluate the difference between the two studies and the corresponding p-value is 0.812, which indicates we could not reject the null hypothesis of no difference between the two studies. In total, we have 15 death events within the two studies. SSSuperPCA outperformed supervised PCA without stability selection again with head and neck cancer data. It increased the Uno’s C-index, log-rank statistics of Supervised PCA from 0.658 to 0.705 and 2.214 to 2.301 respectively. The top three selected stable features for head-and-neck cancer data were zone-size non-uniformity, large zone high gray-level emphasis and small zone emphasis. All the three texture features are the components of gray-level size zone matrix. The three texture features are measures for intratumor heterogeneity and homogeneous. Table 4 shows the average measurements for the six algorithms with head and neck cancer data. Same as the non-small cell lung cancer data set, Cox proportional hazards regression model, lasso, and elastic-net based Cox proportional hazards regression model generated almost identical results with Uno’s C-index of around 0.640, Brier Score equal to 0.062, and log-rank statistics close to 1.05. Unlike the non-small cell lung cancer data set, survival random forests had better survival predictions for head-and-neck cancer when compared to other algorithms except for SSSuperPCA. Figure 2 shows the scatter plots of the SSSuperPCA against the other five algorithms when considering Uno’s C-Index and scatter plots for Brier score, and log-rank statistics are provided in Supplementary Figure 3.

**Table 4.**
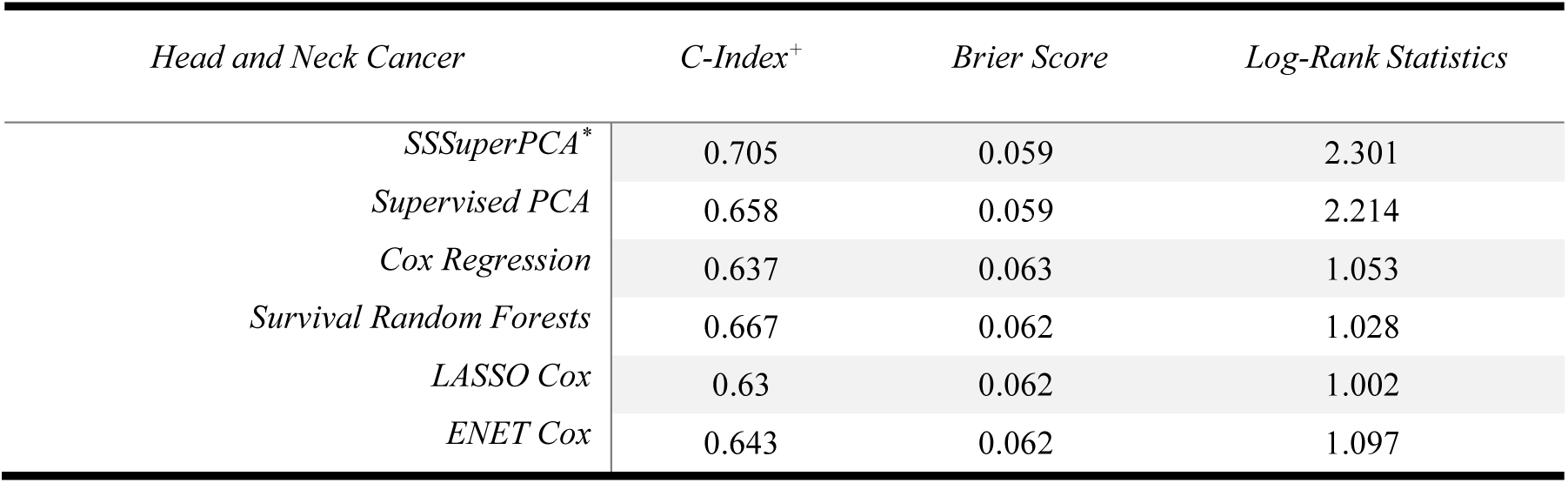
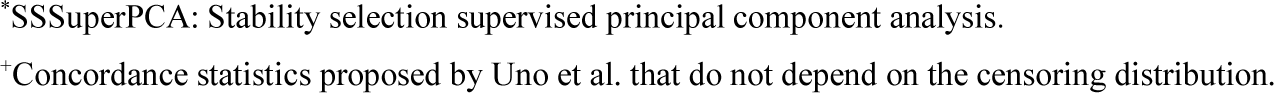
The comparison of stability selection supervised principal component analysis with five machine learning algorithms with data head and neck cancer. The results are the average values that based on 100 times random training/testing splits.

| *Head and Neck Cancer* | *C-Index<sup>+</sup>* | *Brier Score* | *Log-Rank Statistics* |
| --- | --- | --- | --- |
| *SSSuperPCA\** | 0.705 | 0.059 | 2.301 |
| *Supervised PCA* | 0.658 | 0.059 | 2.214 |
| *Cox Regression* | 0.637 | 0.063 | 1.053 |
| *Survival Random Forests* | 0.667 | 0.062 | 1.028 |
| *LASSO Cox* | 0.63 | 0.062 | 1.002 |
| *ENET Cox* | 0.643 | 0.062 | 1.097 |

\*SSSuperPCA: Stability selection supervised principal component analysis.
<sup>+</sup>Concordance statistics proposed by Uno et al. that do not depend on the censoring distribution.

Figure 3 shows the density plots of the two data sets regarding the three measurements, which can provide an overall view of the performance of the six algorithms. Considering the top algorithms compared in Leger’s study, survival regression combined with minimum redundancy maximum relevance would give the best survival prediction for head and neck cancer data with Uno’s C-index, Brier score, log-rank statistics equal to 0.684, 0.061, and 1.413, respectively.

**Figure 3.**
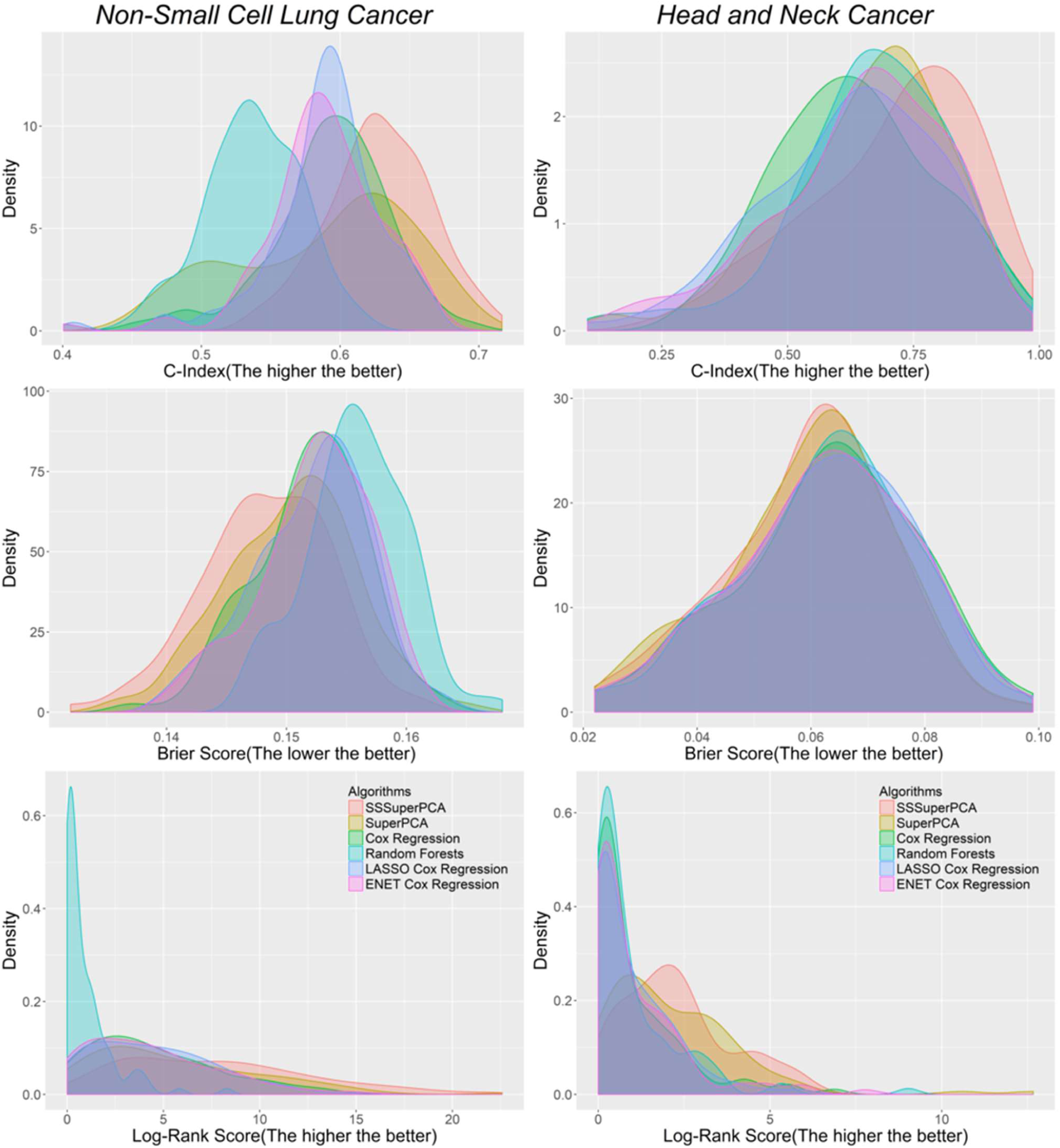
Density plots of three measures of six algorithms for the two data sets.

## IV. DISCUSSION AND CONCLUSION

In this paper, we proposed a new algorithm for radiomics analysis with right censored survival data by coupling boosted stability selection with supervised principal component analysis. This approach allows us to identify a set of stable features that are highly associated with the survival outcomes and predict the survival in a simple yet meaningful manner. Using two different biomedical image cancer data sets, we successfully demonstrated that our algorithm is able to identify a set of biologically meaningful stable features with high predictive ability from the biomedical images. These biologically meaningful features can help researchers or doctors better predict the survival of cancer patients and help doctors in making treatment decisions.

In our experiments, non-small cell lung cancer and head and neck cancer data sets were used to evaluate the performance of SSSuperPCA. Results from both sets of data showed consistent pattern that SSSuperPCA could improve the performance of supervised PCA and also outperformed other regression and machine learning algorithms. In our experiments, we aggregated the stable features that were selected by our algorithm SSSuperPCA in the 100 runs for the two data set and calculated the frequency of these stable features. Two non-texture features that quantify the compactness of tumor shape and one texture feature that assess the homogeneity of tumor characteristics were identified as top three highly selected stable features from the non-small cell lung cancer data. Three texture features, as parts of gray-level size zone matrix, describe the intratumor heterogeneity and heterogeneity were discovered from the head and neck cancer data. These type of radiomics features are also suggested as the potential biomarkers for lung cancer and head and neck cancer in other studies [6, 25]. The grey level non-uniformity and wavelet grey level nonuniformity highpass-lowpass-highpass that measure the intratumor heterogeneity, are two of the four consistency radiomics signatures for the prediction of survival in Aerts’s study. The results demonstrated that our algorithm is able to identify stable features that have higher prognostic ability. The stable features identified from the head and neck cancer data are not as stable as the non-small cell lung cancer data set. This could be due to the smaller sample size or a different disease type with more complex bioimages than those in the other data set. In addition, a simulation study based on non-small cell lung cancer data was also conducted to assess the performance of SSSuperPCA under different scenarios. In general, when the number of stable features increased or the sample size increased, the performance of our algorithm improved.

There are two features selection parts involved in SSSuperPCA. In the first stage, stability selection with boosted C-index is employed to identify a set of stable features that are correlated with the right-censored survival outcome while controlling for the per-family error rate. Unlike some conventional feature selection methods that often fail for high-dimensional data, stability selection with boosted C-index can provide a transparent principle for variable selection and yield consistent results. It is then followed by a semi-supervised strategy through supervised principal component analysis that enables SSSuperPCA to identify the gross correlation structure along with the corresponding survival outcomes and to pare down the influence of these less informative features. With the combined two-staged feature selection, SSSuperPCA can significantly reduce the data dimensionality and identify informative stable features as well. One drawback of our approach is that the selection model used in stability selection and the prediction model used in supervised PCA are based on Cox proportional hazards model. Once the assumption is violated, the prediction results could be misleading, and we may consider replacing the Cox proportional hazards model to accelerated failure time model with a certain distribution, for example, log-normal distribution or Weibull distribution. However, this may not be a severe problem under some circumstances as the hazard ratio could be interpreted as the geometric mean of hazard ratio over time points when the assumption is violated [26]. Although our application data sets are based on CT scans, our approach can be applied to MRI bioimage data as input as well. In conclusion, our approach is able to pick up a set of stable features via boosted survival model from radiomics data, control the per-family error rate and perform well in survival prediction. While the field is still in its infancy, the proposed algorithm would motivate and draw the interest of other researcher to develop novel algorithms for bioimage informatics analysis.

